# Glutamatergic adaptation to stress in medial prefrontal cortex underlies risk and resilience for pessimistic beliefs

**DOI:** 10.1101/2020.06.03.131946

**Authors:** Jessica A. Cooper, Makiah R. Nuutinen, Victoria M. Lawlor, Brittany A. M. DeVries, Elyssa M. Barrick, Shabnam Hossein, Daniel C. Cole, Chelsea V. Leonard, Andrew P. Teer, Grant S. Shields, George M. Slavich, Dost Ongur, J. Eric Jensen, Fei Du, Diego A. Pizzagalli, Michael T. Treadway

## Abstract

Stress is a major risk factor for the development of mental illness, including major depressive disorder (MDD), yet the underlying biological mechanisms remain unclear. Particular challenges lie in disentangling adaptive versus maladaptive responses to repeated stress exposure. Preclinically, stress-induced changes in glutamatergic function have been frequently observed in the medial prefrontal cortex (mPFC), a key region for mediating adaptive stress responses. Here, we examined stress-induced changes in mPFC glutamate using magnetic resonance spectroscopy (MRS) in four human samples varying in perceived stress exposure. Changes in mPFC glutamate following an acute stressor were reliably moderated by recent perceived stress in healthy controls. This adaptive glutamate response was absent in unmedicated individuals with MDD and was associated with excessively pessimistic beliefs as assessed via ecological momentary assessments over a 1-month follow-up period. Taken together, these data provide novel evidence for glutamatergic adaptation to stress in mPFC that is significantly disrupted in MDD.

## Introduction

Stress is a major risk factor for physical and psychological health problems^1,2^, and has been strongly linked to the onset of major depressive disorder (MDD)^3-5^. Although ‘stress’ is often broadly defined, prior research has divided this construct into ‘good stress’, ‘tolerable stress’, and ‘toxic stress’, with the latter being associated with significant risk for physiological damage and mental illness^6,7^. Key features of toxic stress include a lack of predictability and controllability^8^ as well as stressors related to social threat, such as isolation, rejection and exclusion^9,10^. One of the most widely-replicated consequences of stress is stress-induced anhedonia, resulting in behavioral inhibition and a failure to pursue rewards^11-13^. Additionally, elevated perceptions of stress have been found to confer particular risk for blunted reward processing^8,12,14,15^. To date, however, the neural mechanisms of stress-induced anhedonia remain unclear.

The medial prefrontal cortex (mPFC) has emerged as a critical region that may underlie stress-induced anhedonia. A robust preclinical literature has elucidated numerous negative effects of stress in mPFC, including glutamate-mediated excito-toxicity that may result from frequent elevations of circulating glucocorticoids^16,17^. In the rodent mPFC, for example, initial stress exposure has been shown to increase extracellular glutamate^18^, potentiate post-synaptic excitatory currents^19^, and up-regulate surface expression of glutamate alpha-amino-3-hydroxy-5-methyl-4-isoxazolepropionic acid (AMPA) and *N*-methyl-*D*-Aspartate (NMDA) receptors^20^. These effects have, in turn, been linked to adaptive changes, including short-term enhancements in learning and memory. With repeated stress exposure, however, glutamate release in response to subsequent acute stressors shows rapid habituation^18^. Similarly, animals previously exposed to chronic stress demonstrate reduced potentiation of glutamatergic signaling when faced with a subsequent stressor^21,22^. This reduction in mPFC glutamate in response to stress and concomitant reductions in dendritic arbors and spines in mPFC^23-26^ have been proposed as possible protective mechanisms that facilitate a necessary adaptation to repeated toxic stressors^6,7^.

Localization of the above effects to the mPFC is particularly relevant for understanding how perceived stress may lead to the development of stress-related psychopathology. Substantial work has consistently implicated overlapping roles for the mPFC in coordinating behavioral and endocrine responses to stress^27,28^ as well as the valuation of expected rewards^29-31^. The mPFC in particular plays critical roles in representing the expectations and probabilities for future outcomes^32,33^. Additionally, animal studies have strongly implicated this region in both risk and resilience for learned helplessness behavior, where individuals form expectations that their actions are incapable of impacting future outcomes^34^. Taken together, this literature suggests that repeated stress exposure may significantly alter mPFC glutamate function, which in turn may contribute to depressive phenotypes. A critical unanswered question, however, is the extent to which the attenuated mPFC glutamate responses to new stressors represents a protective adaptation or a negative consequence, given individual perceptions of recent stress.

Here, we used magnetic resonance spectroscopy (MRS) to examine changes in glutamate following an acute stressor and how these changes relate to perceived stress. Importantly, we used an acute stress manipulation designed to be unpredictable, minimally controllable and included a strong social-threat component^35^. Glutamate responses to this acute stressor were first evaluated in a sample of healthy adults who varied in levels of recent perceived stress. Based on the preclinical studies described above, we hypothesized that for healthy individuals with low levels of perceived stress, mPFC glutamate would increase following an acute stressor, whereas for individuals with higher levels of perceived stress, mPFC glutamate would decrease. To ensure the stability our results, we then replicated this experiment in a second sample of healthy adults. To determine the specificity of these effects to acute stressor (as opposed to mere exposure to any cognitive task), we next evaluated a third sample of healthy adults using a “no stress” control manipulation designed to mimic the sensory and cognitive components of the acute stressor. Next, we examined the relationship between acute stress-induced mPFC glutamate changes and recent perceived stress in a sample of subjects with MDD, hypothesizing that adaptive mPFC glutamate stress responses would be disrupted in MDD. Finally, to understand how this disrupted response might relate to anhedonia, we used ecological momentary assessment to evaluate associations between mPFC glutamate function and reward processing in daily life.

## Results

### Effects of the Acute Stress Manipulation on Mood and Salivary Cortisol

Participants in this study included healthy controls (HC) across three independent samples and a fourth sample of unmedicated patients meeting criteria for current Major Depressive Disorder (MDD; see **Table 1** for demographic information). After completing interview and self-report measures, all participants completed an MRI scanning session that included two MRS assessments of mPFC metabolites on a 3 Tesla (3T) scanner using a well-validated MRS protocol with excellent test-retest reliability (see Online Methods and **Fig. 1a-c**). In between the first and second MRS scans, participants in two of the healthy control groups and the MDD group completed an acute stress manipulation (Maastricht Acute Stress Task; MAST^35^), whereas the third sample of healthy control participants completed a No-Stress Control (NSC) manipulation.

**Table 1:** Demographics and self-report measures.

|  | Healthy control stress (*n*=25) | Healthy control stress replication (*n*=22) | No stress control (*n*=18) | Major depressive disorder stress (*n*=23) |
| --- | --- | --- | --- | --- |
| Sex (% Female) | 60.00% | 68.80% | 77.80% | 69.60% |
| Race (White/Caucasian) | 76.00% | 59.10% | 38.90% | 73.90% |
| Race (Asian) | 8.00% | 27.30% | 38.90% | 8.70% |
| Race (Black/African American) | 16.00% | 13.60% | 22.20% | 17.40% |
| Age | 26.04 ± 6.20 | 28.36 ± 8.21 | 23.44 ± 4.40 | 29.87 ± 10.61 |
| PSS | 10.12 ± 3.70 | 9.60 ± 5.15 | 11.50 ± 5.79 | 27.91 ± 5.68 |

**Note.** PSS = Perceived Stress Scale

To confirm the success of our stress manipulation, we first examined changes in mood using an adapted version of the visual analogue mood scale (VAMS^36^; see **Online Methods**). Participants were included in a 4 (*Timepoint*) x 3 (*Group*) repeated-measures ANOVA if they had VAMS data from all four timepoints (*N*=77). *Group* included No Stress Control (NSC), Healthy Control Stress (combined samples), and participants with major depressive disorder (MDD). We found a significant main effect of *Timepoint* (*F*_(2.49, 184.56)_ = 8.37, *p* < .001), and main effect of *Group* (*F*_(2,74)_ = 6.04, *p* =.004), as well as a significant *Timepoint* x *Group* interaction (*F*_(4.99,184.56)_ = 3.77, *p* =.003). Among participants who completed the acute stress manipulation, we observed a significant effect of *Timepoint* (*F*_(2.35, 141.16)_ = 11.30, *p* < .001) and main effect of *Diagnostic Group* (*F*_(1,60)_ = 11.01, *p* = .002), but no significant *Timepoint* x *Diagnostic Group* interaction (*F*_(2.35 141.16)_ = 1.25, *p* = .294), indicating that the MDD and control groups exhibited similar decreases in mood following the acute stressor, whereas MDD participants reported higher negative emotional experience overall (**Fig. 1d**). We additionally compared healthy control participants who completed the stressor vs no stress control. While the main effect of *Acute Stress* was not significant (*F*_(1, 54)_ = 0.27, *p* = .606), we observed as significant *Timepoint* x *Acute Stress* interaction (*F*_(2.44,131.53)_ = 7.77, *p* < .001). Whereas healthy control participants who completed the stress manipulation showed peak negative emotional experience following the MAST stressor, negative affect for the NSC group was consistent throughout the scan and lowest at the end of the study (**Fig. 1d**).

In addition to mood effects, we examined changes in salivary cortisol–a widely used marker of the stress response (**Fig. 1e**). We compared cortisol values taken from immediately prior to the onset of the acute stressor (and after habituation to the scanner environment) to the two post-stressor time points collected approximately 20- and 40-minutes post-stressor. All timepoints were scaled relative to the pre-stress timepoint to represent percent change in cortisol from baseline and included in a repeated-measures ANOVA. Participants were only included in this analysis if they had sufficient cortisol from all 3 timepoints (*N*=83). We observed a marginally significant effect of *Timepoint* (*F*_(1.62,129.94)_ = 3.03, *p* = 0.063), marginally significant main effect of *Group* (*F*_(1,80)_ = 2.86, *p* = 0.063), and significant *Timepoint* x *Group* interaction (*F*_(3.25,129.94)_ = 3.19, *p* = 0.023). Among participants (healthy controls and MDD subjects) who completed the stress manipulation, we observed a main effect of *Timepoint* (*F*_(1.57, 98.66)_ = 6.93, *p* = 0.003) and significant quadratic effect of *Timepoint* (*F*_(1,63)_ = 11.49, *p* = 0.001); conversely, the *Timepoint* x *Diagnostic Group* interaction (*F*_(1.57, 98.66)_ = 1.45, *p* = 0.239) and the main effect of *Diagnostic Group* (*F*_(1,63)_ = 1.31, *p* = 0.256) were not significant. Among healthy control subjects, we observed a significant *Timepoint* x *Acute Stress* interaction (*F*_(1.54,90.95)_ = 5.28, *p* = 0.012) and a significant *Timepoint* x *Acute Stress* quadratic contrast (*F*_(1,59)_ = 9.05, *p* = 0.004). Whereas cortisol increased relative to baseline for healthy controls at the first timepoint following the stress manipulation (*t*_42_ = 3.33, *p* = .002), participants in the NSC group showed a slight decrease in cortisol concentration following the no stress control manipulation (*t*_17_ = -2.18, *p* = .044).

**Figure 1:**
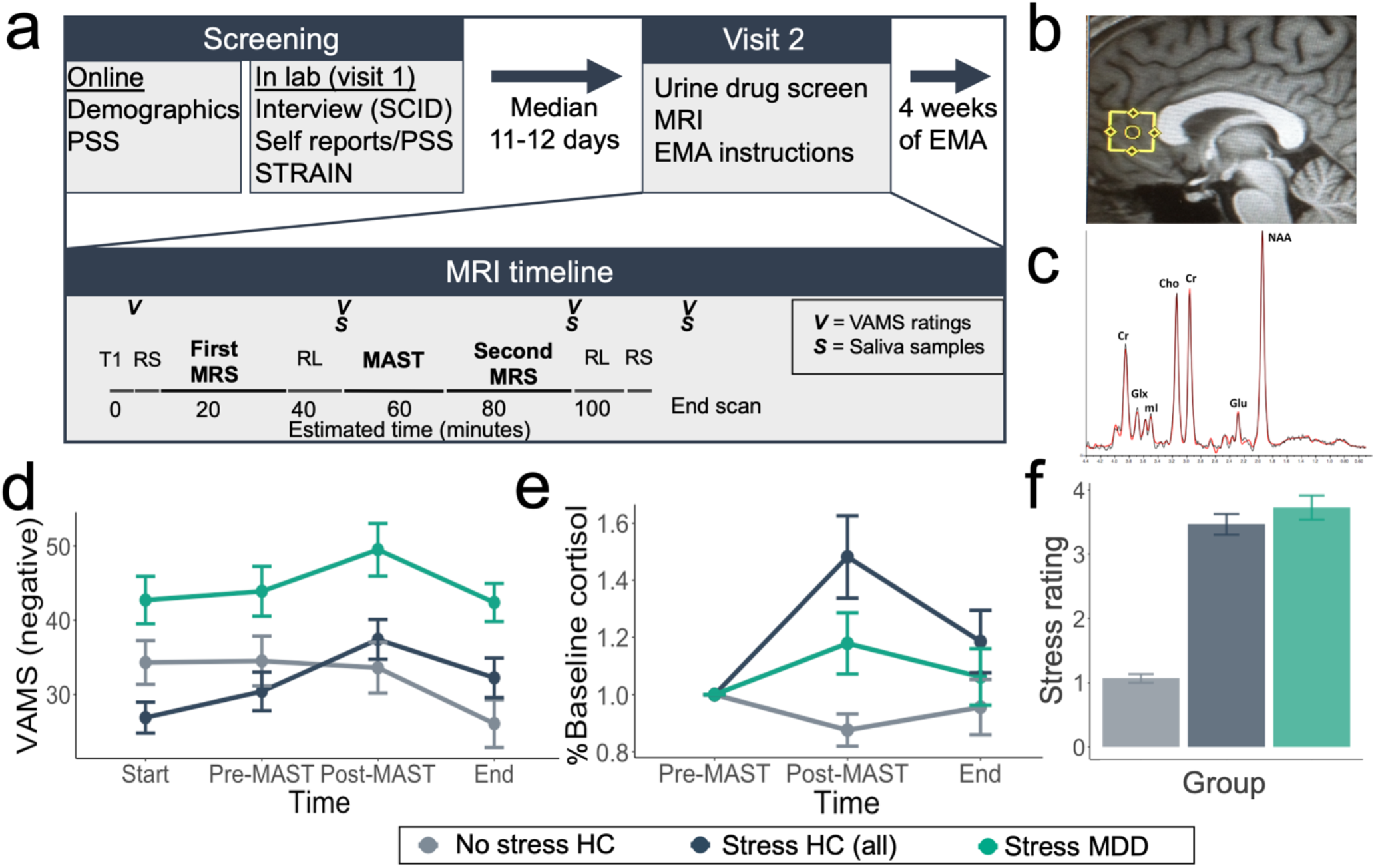
Study design and effects of stress on mood and salivary cortisol. **a)** Schematic diagram of the study visits and timing of MRS, fMRI RL task, VAMS and saliva measurements. Note that the Healthy Control Stress sample did not complete resting state scans, STRAIN, or EMA. EMA=Ecological Momentary Assessment; MRS=Magnetic Resonance Spectroscopy; PSS=Perceived Stress Scale; RL=Reinforcement Learning; RS=Resting State; SCID=Structured Clinical Interview for DSM Disorders; STRAIN=Stress and Adversity Inventory. MAST=Maastricht Acute Stress Test; VAMS=Visual Analog Mood Scales. **b)** Representative MRS voxel placement **c)** Representative MRS spectra (black) and LC model fit (red) with labeled metabolite peaks. Cr = creatine; Glu = glutamate; Glx = glutamine/glutamate/glutathione; Cho = choline; ml = myo-inositol; NAA = N-acetylaspartic acid. **d)** Effect of MAST acute stress task and No Stress Control (NSC) on mood. Items are coded such that higher scores indicate greater negative emotional experience and averaged across items. **e)** Salivary cortisol response to acute stress manipulation and NSC. Graph depicts percent change in salivary cortisol from the time-point immediately prior to the onset of the MAST stressor (Pre-MAST). **f)** Subjective stress ratings for each group (1-5). All error bars are standard error of the mean. NSC=No Stress Control; HC=Healthy Control participants; MDD=participants with Major Depressive Disorder.

Finally, we examined participants’ subjective ratings collected at the end of the scan (*N*=82), which included their subjective levels of stress, unpleasantness, and difficulty of the water/counting manipulation (**Fig. 1f**), using ANOVA, with *Group* (HC stress, NSC, and MDD stress) as a between-subjects factor. Main effects of *Group* were highly significant for all three questions (*ps* < 1.0*10^-13^), driven by lower ratings of the NSC group. For participants who completed the stress manipulation, no significant effects of *Diagnostic Group* were observed for subjective levels of stress, unpleasantness, or difficulty (*p*s>.18).

### Effects of Perceived Stress on mPFC Glutamate following Acute Stress Manipulation in Healthy Control Participants

Having established the validity of our acute stress and NSC manipulations, we next sought to test our primary hypothesis regarding the effects of acute stress on mPFC glutamate in the first Healthy Control Stress sample (*n*=25; McLean Hospital sample). We hypothesized that recent perceived stress as measured by the Perceived Stress Scale (PSS^37^) would predict changes in mPFC glutamate under stress such that healthy individuals with low PSS scores would show greater mPFC glutamate following the acute stress manipulation relative to those with higher PSS scores. Consistent with this hypothesis, percent change in mPFC creatine-normalized glutamate (%ΔGlu; Equation 1) was inversely correlated with participants’ PSS scores (*r*_s_ = -.457, *p* = .022). Individuals with low PSS scores exhibited an increase in mPFC glutamate following acute stress, whereas individuals with higher PSS scores showed either no change or a slight decrease in mPFC glutamate levels (**Fig, 2a**).

To confirm the stability of the relationship between PSS scores and %ΔGlu, we collected a second independent sample of healthy control participants (*n*=22; Healthy Control Stress Replication; Emory sample). As in our first sample, PSS scores were inversely correlated with %ΔGlu with a similar effect size (*r*_s_ = -.517, *p* = .014; **Fig. 2b**). Relationships between %ΔGlx and PSS are reported for both groups in **Supplementary Fig. 3** and were consistent with correlations observed for %ΔGlu. No main effects of acute stress on Glu/Cr or Glx/Cr were observed for either sample (paired-*t p*s > .7; **Fig. 2e,f, Supplementary Fig. 3e,f**), though we note that when looking only at individuals reporting low levels of recent perceived stress (PSS scores <10), the acute stress manipulation did evoke a significant increase in mPFC Glu (*t*23 = 2.39, *p* = 0.026).

**Figure 2:**
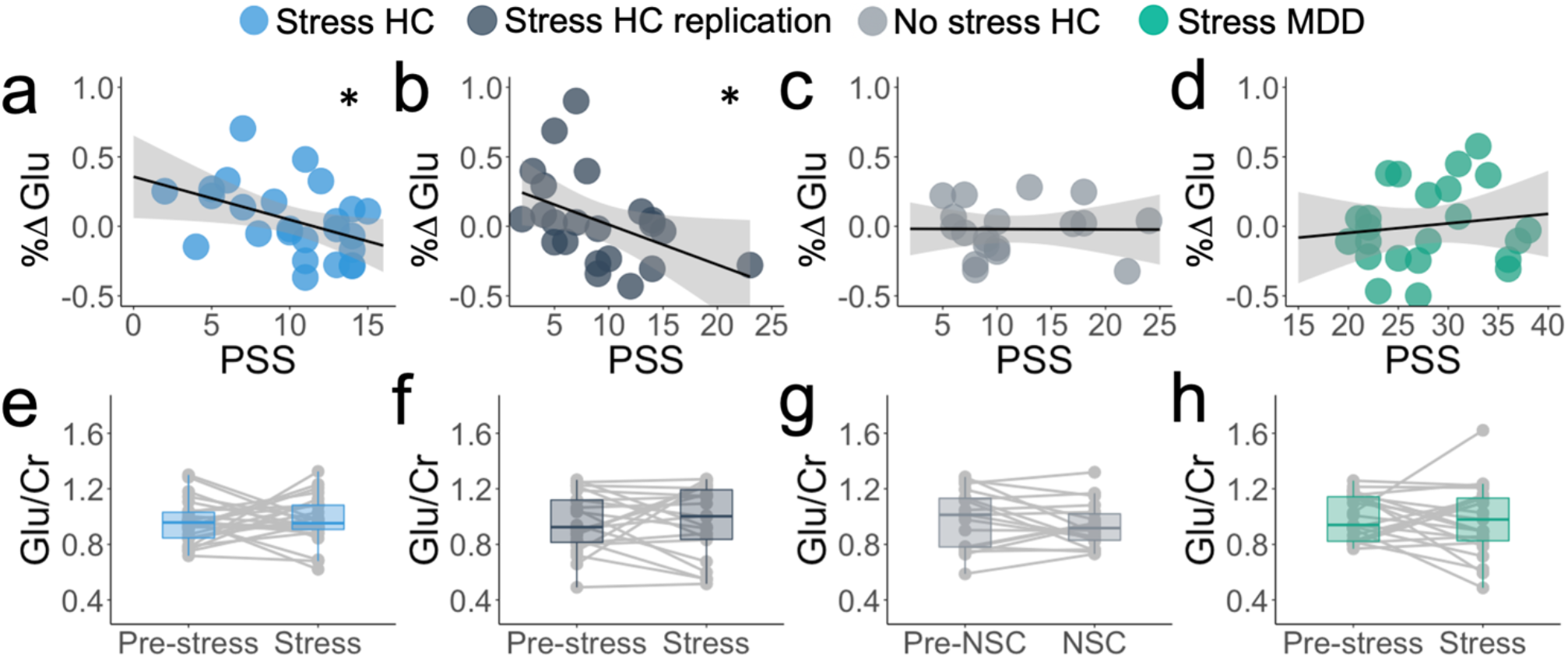
Changes in mPFC glutamate in response to acute and perceived stress **a)** Association between perceived stress (PSS scores) and percent change in Glu signal (%ΔGlu; *r*_s_ = -.457, *p* = .022) in healthy control stress sample. **b)** Association between perceived stress (PSS scores) and percent change in Glu signal (%ΔGlu; *r*_s_ = -.517, *p* = .014) in the healthy control stress replication sample. **c)** Association between perceived stress (PSS scores) and percent change in Glu signal (%ΔGlu; *r*_s_ = -.023, *p* = .928) in no stress control sample. **d)** Association between perceived stress (PSS scores) and percent change in Glu signal (%ΔGlu; *r*_s_ = .126, *p* = .567) in participants with major depressive disorder. Shaded area on a-d represents 95% confidence interval, \**p*<.05. **e-h)** Glu/Cr ratios before and after MAST in healthy control stress sample (e), healthy control stress replication (f), no stress control (g), and subjects with major depression (h).

Next we sought to determine whether the association between PSS scores and %ΔGlu following the acute stress manipulation was specific to the acute stress manipulation as compared to the No Stress Controls (NSC) condition. Participants in the NSC condition showed no association between %ΔGlu and PSS scores (*r_s_=* -.023, *p* = .928; **Fig, 2c**). To confirm that the association between PSS scores and %ΔGlu was significantly stronger during the acute stress manipulation relative to the NSC condition, we additionally examined the interaction between the acute stress manipulation and PSS using hierarchical linear regression. *PSS Score* and dummy-coded *Acute Stress* condition were entered in the first block, whereas *Study Site, Age, Sex*, and the *PSS x Acute Stress* condition interaction term were entered in the second block using stepwise selection. The *PSS x Acute Stress* interaction term and *Age* were both significant predictors of %ΔGlu. The *PSS x Acute Stress* condition interaction term was associated with decreased %ΔGlu (*β* = -.365, *t*_60_ = -2.03, *p* = .047), while *Age* was associated with increased %ΔGlu (*β* = .289, *t*_60_ = 2.39, *p* = .020). No other variables were significant predictors of %ΔGlu (*p*s > .1; see **Supplementary Table 3**). This model explained a significant proportion of the variance in %ΔGlu (adjusted-*R*^2^ = .194; *F*_(4,60)_ = 4.85, *p* = .002), and the change in *R*^2^ from including the *PSS x Acute Stress* interaction was significant (ΔR^2^ *F-change*_(1,60)_ = 4.12, *p* = .047). The model was also run with %ΔGlx and revealed a similar pattern of results (see **Supplementary Table 3**).

Finally, to test whether the effects described above were attributable to a global association with PSS across metabolites, we additionally ran the same regression model to predict %ΔCholine. This model did not explain a significant portion of variance in %ΔCholine (adjusted-*R*^2^ = .015, *F*_(2,62)_ = 1.49, *p* = .233) (See **Fig. 3c** for PSS-metabolite effect sizes).

### Effects of Objective Stress on mPFC Glutamate following Acute Stress Manipulation in Healthy Control Participants

While the Perceived Stress Scale measures how unpredictable and uncontrollable respondents find their life (i.e. stress appraisal), it does not yield an objective assessment of stressors experienced by each participant. To provide a more objective characterization of stress exposure, we used the recently-developed computer-adapted Stress and Adversity Inventory (“STRAIN”^38^), though we note that the STRAIN was not available at the time of data collection for the McLean sample. An advantage of the STRAIN is that it provides a more objective quantification of the number of all stressors experienced, and can therefore help determine whether observed associations with the PSS were more likely driven by stress exposure or perceptions of stress. Interestingly, we found no significant associations between %ΔGlu and STRAIN assessments of count or severity of either acute life events and chronic difficulties (*r* values ranged *from -.148 to .102, ps >.5).* We additionally examined associations between %ΔGlu and STRAIN assessments of count and severity of acute life events and chronic difficulties experienced only within the last year, again finding no significant associations (r values ranged from .008 to .089, *p*s>.4).

Taken together, our results suggest that perceived stress reliably predicts changes in mPFC glutamate following acute stress in healthy individuals, and that our observed associations with PSS may be related to subjective appraisal of recent stress. Moreover, the fact that all participants in these samples had no history of psychiatric illness despite moderate levels of PSS suggests that the observed decrease in mPFC glutamate as PSS scores increased may reflect a beneficial adaptation.

### Effects of Acute and Perceived Stress on Glutamate in Major Depressive Disorder (MDD)

To further understand how perceived stress may be drive mPFC glutamate responses to an acute stressor, we next evaluated a sample of participants with current MDD, a disorder strongly linked to stress exposure and significant elevations in PSS scores^39,40^. In contrast to our healthy control samples, PSS scores and %ΔGlu following the acute stress manipulation were not significantly correlated in participants with MDD (*r*_s_= .126, *p* = .567; **Fig. 2d**). We used hierarchical linear regression across all three samples that completed the acute stress manipulation, with *PSS Scores* and dummy-coded *Diagnostic Group* (HC or MDD) entered in the first block and *Age, Sex, Study Site*, and the *PSS x Diagnostic Group* interaction term entered in the second block using stepwise selection. Only *PSS* (*β* = -.986, *t*66 = -3.10, *p* = .003) and the *PSS x Diagnostic Group* interaction term (*β* = .817, *t*66 = 2.56, *p* =

.013) were significant predictors of %ΔGlu (**Fig. 3a, Supplementary Table 4**). This model explained a significant proportion of the variance in %ΔGlu (adjusted-*R*^2^ = .096, *F*_(3,66)_ = 3.43, *p* = .022) and the change in *R*^2^ from including the *PSS x Diagnostic Group* interaction was significant (ΔR^2^ *F*-change_(1,66)_ = 6.55, *p* = .013). The model was also run with %ΔGlx and revealed a similar pattern of results (**Supplementary Table 4**).

**Figure 3:**
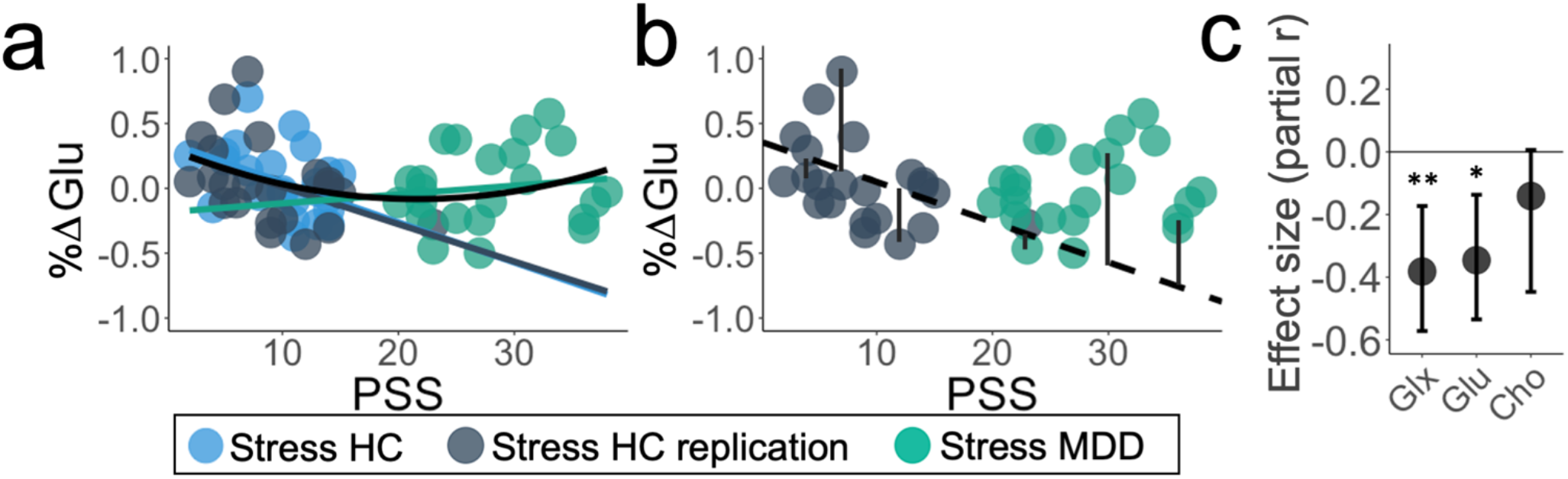
Relationship between stress and metabolites. **a)** Relationship between PSS and %ΔGlu in all participants who completed the acute stress manipulation. Group-level linear trends (colors) and combined quadratic effect (black) are overlaid. **b**) Maladaptive glutamate response was calculated for the healthy control stress replication sample (grey) and participants with major depressive disorder (green), defined as the residual between the observed %ΔGlu and expected %ΔGlu, estimated using the linear function of the healthy control stress sample, shown in black. %ΔGlu expected = .36 - .03*PSS. **c)** Partial effect size (Pearson’s *r*, controlling for age and sex) between PSS and %ΔGlx (Glutamate and Glutamine), %ΔGlu (Glutamate), and %ΔCho (Choline), in all healthy controls who completed the acute stress manipulation. Estimates and error bars (95%CI) were estimated using bootstrapping with 1000 samples. \**p*<.05, \*\**p*<.01.

Although both samples of healthy control participants showed a negative relationship between PSS and %ΔGlu, PSS and %ΔGlu were not significantly correlated in MDD participants. Additionally, without considering diagnostic status, the relationship between PSS and %ΔGlu after being exposed to the acute stressor was predicted by a quadratic function (adjusted-*R*^2^ = .069, *F*_(2,69)_ = 3.55, *p* = .034; **Fig. 3a, Supplementary Table 5**). Relative to the linear function (i.e. only including PSS), the introduction of the quadratic PSS term explained an additional 7.2% of the variance in %ΔGlu (ΔR^2^ *F*-change_(1,67)_ = 5.34, *p* = 024).

### Main Effect of Depression on mPFC Glutamate

The main effect of depression on glutamate signal and interactions with acute stress (**Fig. 2e-h)** were examined using a repeated-measures ANOVA, with levels of glutamate (Glu/Cr) at each *Timepoint* (pre- and post-manipulation) as a within-subjects factor, and *Diagnostic Group* (MDD/control) as a between-subject factor. The main effect of *Diagnostic Group* (*F*_(1,68)_ = .197, *p* =. 658), *Timepoint* (*F*_(1,68)_ = .001, *p* = .975) and *Timepoint x Diagnostic Group* interaction (*F*_(1,68)_ = .150, *p* = .699) were all nonsignificant (see **Supplementary Table 2a-d** for metabolite values). This comparison was also conducted using Glx/Cr, finding consistent results (*p*s > .9). Glu/Cr and Glx/Cr ratios at baseline showed no differences between MDD participants and healthy controls (*p*s > .49), nor were Glu/Cr and Glx/Cr ratios different between the diagnostic groups after being exposed to the acute stress manipulation (*p*s > .9). These results suggest that mPFC glutamate at baseline and in response to acute stress did not differ between healthy control and MDD participants.

### mPFC Glutamate Response and Experience of Reward in Daily Life

Next we sought to determine how altered mPFC glutamate responses might be related to expectations about rewarding events in daily life. Because the interpretation of %ΔGlu depends on the PSS, we developed a novel “maladaptive glutamate response” (MGR) metric that represented the difference between the actual %ΔGlu and the level that would be expected given a participant’s rating of recent perceived stress (**Fig. 3b, Equation 2**). To avoid any non-independence in this analysis, the slope used to calculate the MGR in Equation 2 was defined by the McLean Sample only. We then tested whether the MGR was related to the prediction or experience of reward in daily life collected over a 4-week follow-up period using ecological momentary assessment (EMA) in the Emory samples. Our EMA protocol was designed with particular emphasis on inaccuracy of reward estimation for daily activities. Inaccuracy of reward estimation was quantified as the difference between the experienced reward for an activity and the amount of reward that the participant anticipated experiencing, similar to a reward prediction error under reinforcement learning frameworks.

**Figure 4:**
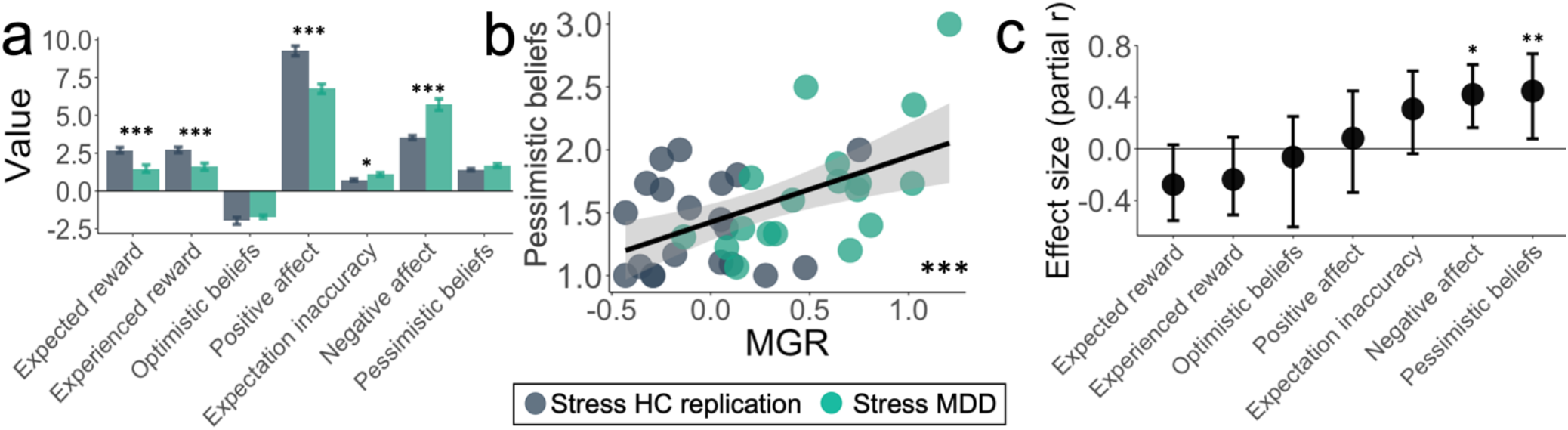
Experience of reward in daily life. **a)** Differences in ecological momentary assessment (EMA) ratings between healthy control (HC) and participants with major depressive disorder (MDD). **b)** Association between maladaptive glutamate response (MGR) and overly pessimistic beliefs from EMA, (*r* = .515, *p* < .001). **c)** Effect sizes (partial Pearson’s *r*) between MGR and EMA variables, controlling for age, sex, and diagnostic group. Note that the optimistic beliefs comparison had one less degree of freedom than other variables, as one participant had no surveys with optimistic beliefs. Effect sizes and error bars (95%CI) were estimated using bootstrapping with 1000 samples and estimated separately for variables with different numbers of observations. \**p*<.05, *\*\*p<.*01, \*\*\**p*<.001.

Compared to healthy controls, participants with MDD reported higher average negative affect (*t*_36_ = 5.62, *p* < .001), lower positive affect (*t*_36_ = -5.44, *p* < .001), lower expected reward (*t*_36_ = -3.98, *p* < .001), and lower experience of reward (*t*_35_ = -3.74, *p* = .001). On average, participants with depression had a lower proportion of responses with accurate prediction (M=.38) than healthy control participants (M=.53; *t*_36_ = -2.35, *p* = .024; see **Supplementary Fig. 4** for distributions). The average magnitude of prediction inaccuracy was also greater in participants with depression (*t*_36_ = 2.45, *p* = .019), indicating less accurate estimations of reward (**Fig. 4a**). We examined directionality of expectation inaccuracies by calculating the mean inaccuracy when expectations were lower than the experienced outcome (“pessimistic beliefs”) and when expectations were higher than the experienced outcome (“optimistic beliefs”). Participants with depression had a higher proportion of responses with pessimistic beliefs (M=.38) than healthy control participants (M=.27; *t*_36_ = 2.11, *p* = .042; **Supplementary Fig. 4**), and marginally higher proportion of trials with optimistic beliefs (M=.24) than healthy control participants (M=.19; *t*_36_ = 1.71, *p* = .095; **Supplementary Fig. 4**). Participants with MDD had marginally higher magnitude of inaccuracies from pessimistic beliefs (*t*_36_ = 1.89, *p* = .066), but did not differ in mean inaccuracies from optimistic beliefs (*p* = .34), suggesting that participants with MDD experienced pessimistic beliefs more often than healthy controls, and that their pessimistic beliefs were more negative than those experienced by healthy controls.

Maladaptive glutamate response (MGR) was compared to EMA variables, controlling for age, sex, and diagnostic group with pairwise exclusions. MGR was positively associated with negative affect (*r*-partial = .422, *p* = .012), as well as pessimistic beliefs (partial-*r* = .461, *p* = .005; **Fig. 4b**), but not optimistic beliefs (partial-r = -.088, *p* = .619; see **Fig. 4c**). When additionally controlling for participants’ PSS scores, the association between MGR and pessimistic beliefs remained significant (*r*-partial = .354, *p* = .040), whereas the relation between MGR and negative affect was marginally significant (*r*-partial = .313, *p* = .071). The relationship between MGR and pessimistic beliefs remained significant when additionally controlling for frequency of pessimistic beliefs (partial-*r* = .399, *p* = .021). Among participants in which both optimistic and pessimistic beliefs were observed (*n*=37) we additionally compared the partial correlations, controlling for age, sex, and diagnostic group, between MGR and pessimistic beliefs and MGR and optimistic beliefs using Steiger’s-Z^41,42^ and found that the relationship between MRG and pessimistic beliefs (*r*-partial = .479) was stronger than the relationship between MGR and optimistic beliefs (*r*-partial = -.064), Z = 2.06, *p* = 0.039.

## Discussion

In this study, we characterized a novel adaptation of mPFC glutamatergic response to acute stress in humans. In two independent samples, we found that healthy individuals exhibited a clear reduction of mPFC %ΔGlu response to a new stressor as their levels of recent perceived stress increased. Critically, this effect was absent for unmedicated individuals with current MDD, suggesting that this absence of adaptation may be a contributor to stress-related psychiatric disease. Further, the extent to which individuals failed to exhibit an attenuated mPFC %ΔGlu response was linked to negative functioning in daily life.

The observed relationship between perceived stress and mPFC Glu suggests an important adaptation to stress among healthy control participants. Importantly, participants from these two samples were confirmed to have no history of psychiatric illness, despite the fact that their perceived stress levels extended into the moderate range^43^. This suggests that some of our participants exhibited resiliency in the face of mild-to-moderate perceived stressors, supporting the notion that attenuated glutamate response may represent an appropriate adaptation to an elevated allostatic load. Under models of allostatic regulation, biological and behavioral responses to an acute stressor should be influenced by levels of recent perceived stress^6,7,16^. Consistent with this framework, preclinical studies have provided clear evidence that glutamate transmission is potentiated by acute stress and stress hormone exposure^17,19,20,44^, and that this effect is reversed if an acute stressor is experienced in the context of recent stress^16,18,21^. Critically, attenuation of the glutamate response among individuals with high perceived stress was attributable to the experience of an acute stressor; when healthy control participants completed a task that matched the cognitive and sensory components of the MAST but was reported as non-stressful (see **Fig. 1d-f**), there was no association between perceived stress and %ΔGlu.

To further determine whether the association between glutamate and perceived stress was adaptive, we next examined the response to acute stress in participants with MDD, a population known to be associated with excessive stress exposure and impaired coping. Participants with MDD reported significantly higher levels of perceived stress relative to healthy control participants. Critically however, these elevated PSS scores did not show the same association with mPFC %ΔGlu following acute stress that was observed in two independent healthy control samples. Indeed, moderation analysis confirmed a significant difference in how PSS scores predicted mPFC %ΔGlu response to acute stress as a function of MDD.

Although MDD participants as a whole did not show the same inverse association between perceived stress and %ΔGlu, we observed significant variability in glutamate change following acute stress. Using the slope from one of the healthy control samples as an independent quantitative estimate for appropriate %ΔGlu given a certain level of perceived stress, we were able to quantify a “maladaptive” %ΔGlu response” (MGR) and examine whether this metric was associated with experience of reward in daily life. During our 4-week EMA follow-up period, we found that the MGR predicted a consistent pattern of inaccurately low expectations for future events—when activities went better than expected, high MGR was associated with reduced accuracy (i.e., activities were expected to be *much less* positive than they actually were). This effect remained strong even while controlling for depression diagnosis, PSS score, and frequency of pessimistic beliefs. It is further notable that this effect was only evident for occasions when events were *better* than expected, suggesting it may play a critical role in stress-induced anticipatory anhedonia. Moreover, our results underscore the critical role for anticipation and expectation setting in the clinical phenomenology of anhedonia^8,45^, as well as the general tendency for depressed patients to make overly negative predictions about future events^46,47^. This interpretation is also consistent with a much broader preclinical and human neuroimaging literature that has repeatedly implicated mPFC as a region involved in estimating the expected value for future options^31,48,49^ and self-related valuation in general^50^.

It is also notable that while a replicable association emerged between mPFC %ΔGlu and the PSS, we did not observe any association between mPFC %ΔGlu and various indices of stress exposure as indexed by the STRAIN. One potential reason for this lack of effect is a difference in timescales between the STRAIN and PSS; while the PSS focuses on appraisal of stressful experiences over the last month only, the STRAIN assesses lifetime stress exposure. That said, we did not observe any significant associations between the STRAIN and glutamate even when limiting measures of stress exposure to the previous year, suggesting that mPFC %ΔGlu in response to stress may be related to stress exposure over even shorter timescales. Alternatively, this dissociation could be attributable to fundamentally different components of stress captured by the PSS and STRAIN. While the STRAIN focuses on more objective quantification of stressful life events, the PSS probes feelings of uncontrollability, unpredictability, and generally feeling “stressed”. This appraisal may be more akin to chronic low-level stressors, feelings of being “stressed out”, and inability to cope that contribute to allostatic load^6,51^. Both explanations are plausible and they are not mutually exclusive. We also note that we were only able to collect the STRAIN in the Emory samples, which may have limited our ability to detect associations. Future research will be needed to determine how adaptation of mPFC %ΔGlu is related to the perception and timescale of stressful experiences.

Collectively, these findings have a number of implications for our conceptualization of biological adaptations to stress and their potential role in psychiatric disorders. Our study reveals that recent perceived stress reliably moderates mPFC glutamate responses to a novel acute stressor in psychiatrically healthy individuals, but not in those experiencing depression. This is notable, as it suggests that the effects of mPFC glutamate levels depend critically on context. Although glutamate dysfunction has been implicated in MDD^16,52,53^, cross-sectional comparisons of resting metabolites may be insufficient to serve as a reliable biomarker. Indeed, a recent metaanalysis of case-control MRS studies in mPFC found no evidence for consistent basal differences in glutamate associated with MDD^54^. Consistent with this result, the present study also did not observe any group differences in baseline glutamate or Glx levels, or in %ΔGlu following acute stress.

The present study is not without limitations. First, our hypotheses regarding stress and mPFC glutamate were primarily informed on the basis of preclinical studies that were able to measure synaptic glutamate and post-synaptic excitatory currents. In contrast, the vast majority of the MRS glutamate signal is driven by intracellular glutamate, and MRS measures of glutamate metabolite concentration cannot be used to make direct inferences about glutamate transmission or synaptic release. However, prior fMRS studies and meta-analyses suggest that pain or stressful stimuli can induce reliable changes in MRS metabolites that are consistent with expected changes based on preclinical studies^55-59^, and our acute stress manipulation did evoke a significant increase in glutamate for healthy individuals with low levels of recent perceived stress. This suggests that while the precise interpretation of changes in glutamate is unclear, it can still serve as a potential biomarker for individual differences in response to stress. A second limitation was the inclusion of only moderate sample sizes, which were partly due to the exclusion of participants with poor-quality MRS data. To address this concern, we recruited a replication sample of healthy controls, and found very similar effect sizes for the relationship between the PSS and %ΔGlu. An additional limitation was the lack of association between %ΔGlu and salivary cortisol responses to stress. Preclinical studies have suggested that glucocorticoids play a critical role in shaping prefrontal glutamate responses to stress^17^, and the lack of association between these variables in our study was unexpected. This may be due to limitations in the temporal resolution of our MRS and saliva measurements. Finally, it should be noted that we were unable to recruit healthy control participants and participants with MDD with fully overlapping distributions of PSS scores, despite a robust pre-screen effort using online recruitment tools. This was not entirely unexpected, as PSS scores are known to be much higher in MDD samples^39,40^; however, it does limit our ability to determine whether the maladaptive glutamate response we observed was driven primarily by the high severity of perceived stress in MDD, the presence of their current depression, or both.

In sum, this study is the first that we know of to identify attenuation of mPFC %ΔGlu as an adaptive response to acute stress in the context of perceived stress, and to demonstrate how this response is impaired in individuals with depression and also associated with overly pessimistic beliefs in daily life. These results advance our understanding of the neurobiological adaptation to stress, and may play a valuable role in identifying new treatment targets and markers of treatment response in human stress-related illness.

## Methods

Methods and any associated references are available in the online version of the paper.

## Online Methods

### Participants

Adults (age 18-60) participated in this study across three independent samples of healthy controls (HC) and a fourth sample of unmedicated patients meeting criteria for current Major Depressive Disorder (MDD). The first sample of healthy control participants (Healthy Control Stress) was recruited at the McLean Imaging Center (McLean Hospital) in response to community advertisements in Boston, MA, whereas the three replication, extension, and patient samples (i.e., Healthy Control Stress Replication, No Stress Control, Major Depressive Disorder Stress) were recruited at the Facility for Education and Research in Neuroscience (FERN) neuroimaging center at Emory University in Atlanta, GA. To ensure a range of perceived stress scores, individuals recruited for the three samples collected at Emory first completed an online eligibility screening in REDCap^60^ that included the Perceived Stress Scale (PSS^37^) and additional demographic and eligibility questions. *Eligibility Criteria:* For healthy control subjects in all samples, participants were excluded for any current or past psychiatric disorder, with the exception of specific phobia, or past alcohol abuse, as assessed by the Structured Clinical Interview for the DSM-IV (SCID)^61^ administered by a trained master’s level clinician. For participants in the MDD group, diagnosis of MDD was confirmed using the SCID. Additional exclusion criteria for participants with MDD included current substance abuse or dependence, obsessive-compulsive disorder, bipolar disorder, active suicidal ideation as assessed by the Columbia-Suicide Severity Rating Scale (C-SSRS^62^, or any form of psychotic disorder. Participants with MDD with comorbid anxiety disorders or post-traumatic stress disorder were not excluded from the study. Participants in all samples were excluded for recent use of illegal drugs or any psychotropic medications, which was confirmed using a urine drug screen immediately prior to scanning.

In total, 124 participants met inclusion criteria and participated in the MRI visit (n_control_= 93, nMDD = 31). Thirteen participants did not finish the scan visit due to time constraints, undiagnosed claustrophobia, subject illness, inability to fit comfortably in the scanner, or scanner malfunction. Exclusion criteria for MRS data included signal to noise ratio (SNR) less than 9, full width at half maximum (FWHM) greater than 0.15, Cramer-Rao standard deviation for glutamate greater than 20, or poor spectral quality based on visual inspection. Quality of MRS data from the McLean sample was reviewed by JEJ, while quality of MRS data from the Emory samples were reviewed by MTT and FD (MR Physicist, blind to study results), with excellent agreement between ratings, Cohen’s κ = .871, *p* < .001. In cases of disagreement, judgement was deferred to FD. Twenty-two participants had at least one MRS session of insufficient quality. One additional participant was excluded for a change in glutamate over 3 standard deviations from the mean, resulting in a final sample size of 88. Only participants from the final sample (“study completers”) were included in subsequent analyses. Sample demographics for study completers in each group are provided in **Table 1** and comorbidities for study completers with MDD are included in **Supplementary Table 1**.

### Study Description

All recruitment and testing procedures were approved by the Partners Institutional Review Board (McLean Hospital) and the Emory University Institutional Review Board. During an initial study visit and after informed consent, participants were interviewed using the DSM-IV SCID^61^ to confirm eligibility criteria and completed self-report questionnaires. During the second visit, participants completed an initial MRS scan, a reinforcement learning (RL) task, and an acute stress or no stress control task (described below), followed by a secondary MRS scan and RL task. Resting-state and task fMRI data were also collected but were not included in these analyses. Salivary cortisol samples were collected before and after the stress (or no stress) manipulation to determine the presence of a stress response (see **Fig. 1a and 1e)**.

### Acute Stress Manipulation

To induce stress during the scanning session, participants completed the Maastricht Acute Stress Task^35^. The MAST is a laboratory stress paradigm that combines alternating periods of well-validated stress-inducing procedures, specifically a cold pressor and performance of serial subtraction in front of evaluators. During the cold pressor, participants were instructed to immerse their hand up to and including the wrist into ice water (1–8°C). Water immersion occurred 5 times for varying time intervals of (30s-90s) using a fixed randomized sequence that was unknown to participants so as to create a sense of unpredictability. Between water immersion periods, participants were asked to perform serial subtraction starting from 2043 and counting down by 17; with every mistake, a neutral evaluator instructed the participant to re-start from 2043. There were 4 serial subtraction blocks, varying in duration between 30s-90s. Although the MAST protocol we followed was not originally developed for the scanner environment, all procedures were completed while the participant remained in position in the scanner. The scanner bed was moved out part way to facilitate access of the participant’s hand to a container of cold water. We note that this protocol represented our own MRI-related adaptation of the MAST, and is slightly distinct from the fMRI adaptation developed by Smeets and colleagues (the “iMAST”^63^) though both procedures are highly similar to the original MAST protocol.

### No Stress Control Manipulation

Participants in the “no stress control” (NSC) condition were instructed to complete a task that followed the same design and timing as the MAST, but used water at a comfortable temperature (26–36°C) instead of cold water and were asked to count aloud starting from one instead of serial subtraction. Frequency and duration of immersion and counting were determined by computer in the same manner as the MAST. This manipulation was designed to be as similar to the MAST stressor as possible without inducing a stress response.

### Salivary Cortisol Analysis

Salivary cortisol was collected as indicated in **Fig.1a**. Samples were stored -20 °C until they were assayed in duplicate for cortisol using a commercially available chemiluminescence immuno assay (CLIA) from IBL-International, Hamburg, Germany (Cortisol Luminescence Immunoassay). Cortisol from saliva samples were assayed at the Laboratory for Biological Health Psychology at Brandeis University (Directors: Dr. Nicolas Rohleder and Dr. Jutta Wolf). Inter- and intra-assay coefficients were below 10%. Changes in salivary cortisol following the MAST are shown in **Fig. 1e**.

### Self-Report Ratings Questionnaires

To assess perceptions of stress, participants were administered the Perceived Stress Scale (PSS^37^). The PSS is a 10-item questionnaire that asks participants about their perceptions of stress over the last month. Importantly, prior studies have shown that this measure predicts individual differences in mPFC responses to reward information^15^ and reward learning abilities^12^, as well as responses to acute stress^64^. Participants in the replication and extension groups also completed the Stress and Adversity Inventory for Adults (STRAIN^38^). The STRAIN is an online stress assessment interview that measures cumulative lifetime exposure to different types of stress and that has been shown to predict numerous health-related outcomes, including self-reported mental and physical health problems^65^ and biological reactivity to acute stress^66^. Variables extracted from the STRAIN included the STRAIN’s two main stressor exposure outcomes (i.e., lifetime stressor count and severity) and indices indicating the specific types of stressors experienced (i.e. count and severity of both acute life events and chronic difficulties).

To measure affective responses to the acute stress paradigm (described below), all participants completed mood ratings using an adapted version of the visual analogue mood scale (VAMS^36^). This scale presents participants with five horizontal lines, each representing a bipolar dimensional mood state: Happy-Sad, Relaxed-Tense, Friendly-Hostile, Sociable-Withdrawn, Quick Witted-Mentally Slow. Participants were instructed to move a cursor on each line to the point that best described their current mood state. This VAMS scale was administered before and after the MAST acute stress manipulation (see **Fig. 1a**). All VAMS ratings were scaled so that higher scores indicated greater negative emotional experience and averaged for each subject to represent negative emotional experience for each timepoint. Changes in VAMS average ratings for study completers are shown in **Fig. 1d**. Following the completion of the MRI scan, participants were asked to rate the stress (or no stress) manipulation on difficulty, stress (**Fig. 1f)**, and unpleasantness on a scale from 1 (Not at All) to 5 (Extremely).

### MRS Acquisition

For both the McLean and Emory sites, MRS data were collected on a 3T Siemens Tim TRIO using a 32-channel phased-array design RF head coil operating at 123 MHz for proton imaging and spectroscopy using an identical Proton MRS sequence developed by JEJ. High-resolution T1-weighted anatomical images were used to position a single 2×2×2 cm voxel in the mPFC such that the posterior edge of the voxel was placed directly in front of the anterior edge of the corpus callosum (see **Fig. 1b** for a representative placement). Proton MRS employed a modified J-resolved PRESS protocol (2D-JPRESS), which collects PRESS MRS spectra at incremental echo-times (TE) to sample the J-resolved periodicity of coupled metabolites (e.g., Glu and Gln) for better spectral resolution^67-69^. Shimming of the magnetic field within the prescribed voxel was done automatically using an automated shimming routine followed by a manual shim to further minimize unsuppressed water linewidth and optimize voxel field homogeneity. Following the additional automated optimization of water suppression power, carrier-frequency, tip angles and coil tuning, the 2D-JPRESS sequence collected 22 echo time (TE)-stepped spectra with the echo-time ranging from 30ms to 350ms in 15ms increments. Acquisition parameters were: repetition time (TR)=2 s, f1 acquisition bandwidth=67 Hz, spectral bandwidth=2 kHz, readout duration=512 ms, NEX=16/TE-step, total scan duration=12 min. The identical sequence was performed twice (Pre-MAST, Post-MAST).

### Test-Retest Reliability of MRS Sequence

The test-retest reliability for J-resolved MRS scans using this protocol at McLean Hospital in an overlapping rostral anterior cingulate cortex ROI has been previously established, with less than 10% variance for Glutamate/Creatine (Glu/Cr) ratios^70^ and intraclass correlation coefficient (ICC) of 0.803^71^. To confirm that we were able to achieve a comparable level of test-retest reliability at the Emory scanning site, six additional subjects completed two consecutive MRS scans using our 2D-JPRESS protocol. Consistent with prior work^71^, ICC values for Glu/Cr metabolites were calculated in SPSS v25 (IBM, Armonk, NY) using two-way mixed models with absolute agreement, finding excellent test-retest reliability (ICC = 0.89, *p* = .017; **Supplementary Fig. 1**).

### MRS Analysis

All spectroscopic data processing and analyses were performed on a Linux workstation. To quantify glutamate (Glu) with the JPRESS data, the 22 TE-stepped free-induction decay (FIDs) were first zero-filled out to 64 points (TE-stepped dimension), Gaussian-filtered, and Fourier transformed. Consistent with validated methods^69,72^, every J-resolved spectral extraction within a bandwidth of 67 Hz was fitted with LCModel^73,74^ and its theoretically-correct template, which used an optimized GAMMA simulated J-resolved basis sets modeled for 2.89 T (the actual field strengths of Siemens Tim Trio scanners)^69,72^. The integrated area under the entire 2D surface for each metabolite was calculated by summing the raw peak areas across all 64 J-resolved extractions for each metabolite. Glu metabolites were expressed as ratios to total creatine (Cr). A representative spectrum and associated LCmodel fit is shown in **Fig. 1c**. %ΔGlu was calculated using Equation 1:

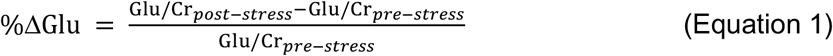

%ΔGlx (Glutamate and Glutamine) and %ΔCholine were also calculated using Equation 1. Creatine ratios for each sample of participants are included in **Supplementary Tables 2a-d**.

### Ecological Momentary Assessment (EMA)

Participants in the replication and extension samples were invited to participate in a four-week ecological momentary assessment (EMA) protocol to assess reward expectation and experience in daily life. EMA data were collected using Qualtrics survey software (Qualtrics, Provo, UT), with survey links sent to participants’ phones via scheduled text messages. Surveys were sent every other day for a period of four weeks. On active survey days, participants received six surveys spaced by two hours. Participants were asked to rate their current affect, indicate their planned activity in the next two hours, indicate whether or not their last planned activity occurred (to provide ratings regarding the outcome of completed activities), and to rate their expected affect. Ratings of current and future affect were collected on a 5-point scale from 1 (“not at all”) to 5 (“extremely”) for positive affect items (enthusiastic, cheerful, and relaxed) and negative affect items (anxious, sad, irritable). Expectations for activities were rated on a 9-point scale from -4 (very negative) to +4 (very positive). For the full question and survey flow, see **Supplementary Fig. 2**. Forty participants completed the EMA protocol (HC: 21, MDD: 19). Two participants were excluded from analysis for having less than 20 usable survey data points, resulting in data from 20 healthy controls and 18 individuals with MDD. Surveys were excluded if they were incomplete, extended beyond the sixth survey of the day, were completed in less than 30 seconds or more than 24 hours, or were not completed within 1 to 3 hours following the prior survey, resulting in usable data from 1,236 surveys from healthy control participants and 1,138 surveys from MDD participants. Inaccuracy of reward estimation was quantified as the difference between the experienced reward for an activity and the amount of reward that the participant anticipated experiencing, similar to a reward prediction error under reinforcement learning frameworks, and included surveys after the first survey of the day (HC = 813, MDD = 659).

### Maladaptive Glutamate Response to Stress

Adaptive glutamate response under stress was characterized using the linear function between recent perceived stress (PSS) and %ΔGlu in the Healthy Control Stress sample. This out-of-sample linear function was used to estimate the expected %ΔGlu (%ΔGlu_exp_) for each participant from the replication sample and patient samples. Maladaptive glutamate response (MGR) was estimated as the difference between observed glutamate change under stress (%ΔGlu_obs_) and %ΔGlu_exp_ (Equation 2), where positive values indicate that mPFC glutamate increased more than expected given the participant’s recent perceived stress, and negative values indicate that change in glutamate was less than expected given recent perceived stress.

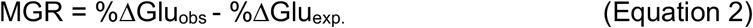

### Statistical Analysis

Change in self-report ratings, task performance, and salivary cortisol were analyzed using separate repeated measures ANOVAs. For cases that violated the sphericity assumption, a Greenhouse-Geisser correction was used. For single sample correlations, Spearman correlations were used to control for possible violations of parametric assumptions that can occur in modest sample sizes and are indicated as r_s_. Interactions between recent perceived stress and the acute stress manipulation in predicting mPFC glutamate levels were examined using hierarchical linear regression, using mean-centered continuous independent variables. Analyses were performed using Matlab 2013B (Mathworks, Natick, MA), SPSS v25 (IBM, Armonk, NY), STATA SE/14 (StataCorp, CollegeStation, TX), and R (RStudio, Boston, MA). EMA data were analyzed using Jupyter Notebooks in Python^75^.

## ACKNOWLEDGEMENTS

This work was supported by the National Institutes of Mental Health R00MH102355 and R01MH108605 to MTT, R37 MH068376 and R01 MH101521 to DAP. JAC was supported by F32 MH115692, DO was supported by K24 MH104449 and GMS was supported by K08 MH103443. Data management through REDCap was supported by UL1 TR000424. The authors also gratefully acknowledge support from Emma Hahn, Nadia Irfan, Asim Lal, Nancy Brooks, Sandra Goulding, Amanda Shamblaw, Amanda Arulpragasam, Nicolas Rohleder, and Laurie Scott. We would like to acknowledge the important contribution of Dr. J. Eric Jensen, who sadly passed away in August of 2017. Through his expertise in MRS physics, Dr. Jensen spearheaded the MRS protocol used in the current studies. It is to Dr. Jensen that we wish to dedicate this manuscript.

## AUTHOR CONTRIBUTIONS

MTT, DAP, DO and JEJ designed the study. JAC, MRN, VML, BAMD, DAP and MTT analyzed the data. JAC, MRN, BAMD, EMB, DCC, CVL, APT and MTT collected data. GSS and GMS provided and analyzed the STRAIN. JAC, VML, SH, and MTT contributed to EMA design and analysis. JEJ developed the MRS sequence and analysis pipeline. DO, JEJ, MTT and FD interpreted MRS data and assessed MRS data quality. JAC, MTT and DAP drafted the manuscript. All of the authors edited and approved the manuscript.

## DATA AVAILABLITY

Note: Any Supplementary Information and Source Data files will be made available upon publication.

