## Supplementary Information for "Glutamatergic adaptation to stress in medial prefrontal cortex underlies risk and resilience for pessimistic beliefs"

**Supplementary Tables: 7**
**Supplementary Figures: 4**

### Supplementary Results

*Demographic effects and effects of birth control on %ΔGlu*

In control analyses, we additionally evaluated the putative effects of demographic variables on %ΔGlu and their potential moderation of perceived stress effects in healthy controls who completed the stress manipulation (combined Emory and McLean samples). We did not observe a significant effect of *Sex* ($p$=.842), nor did we observe a significant *Sex x PSS* interaction ($p$=.355; **Supplementary Table 6**). Change in glutamate was not significantly different between male and female participants ($t_{45}$ = -1.05, $p$=.30). Increasing age was associated with increased %ΔGlu in response to stress ($\beta$ = .273, $p$ = .049), however, we did not observe a significant *Age x PSS* interaction (**Supplementary Table 6**). Among healthy control female participants who completed the stress manipulation, birth control (oral/IUD) was not associated with %ΔGlu under stress ($p$=.647), and did not moderate effects of PSS on %ΔGlu ($p$=0.603; **Supplementary Table 7**)

*Associations between glutamate, cortisol, and mood*

Among healthy controls who completed the stress manipulation, PSS scores were not significantly correlated with cortisol (percentage relative to baseline) at 20- or 40-minutes post-stress, VAMS response (percentage change from T1 to T3), or post-scan subjective ratings, ($p$s>.14). VAMS response and cortisol response at 20 minutes and 40 minutes post-stressor were also not correlated ($p$s>.7). %ΔGlu was not significantly correlated with VAMS change or cortisol response ($p$s>.14). We repeated these comparisons in MDD subjects, finding no significant associations ($p$s>.12).

**Supplementary Table 1: Comorbidities in participants with major depressive disorder.**

| | |
|---|---|
| Anxiety disorders | |
| Generalized anxiety disorder (current) | 8 |
| Panic disorder (current) | 1 |
| Social phobia (current) | 5 |
| Social phobia (past) | 1 |
| Agoraphobia | 1 |
| Substance use and dependence | |
| Past alcohol abuse | 4 |
| Past alcohol dependence | 2 |
| Past THC dependence | 1 |
| Post-traumatic stress disorder (PTSD) | |
| PTSD (current) | 1 |
| PTSD (past) | 3 |

***Note.*** The number of study completers meeting diagnostic criteria for each comorbidity is shown. Participants are included in the count for every disorder for which they met criteria. All participants were free from psychotropic medications. THC = Tetrahydrocannabinol.

**Supplementary Table 2a: Mean Cr ratios and Cramer-Rao standard deviations for MRS metabolites pre-stressor and post-stressor in the healthy control stress sample.**

| Aspartate | | | | | | | | |
|---|---|---|---|---|---|---|---|---|
| Pre-stressor | | | | Post-Stressor | | | | Paired t-test p-value |
| Mean ratio (Cr) | SD ratio (Cr) | Mean Cramer-Rao | SD Cramer-Rao | Mean ratio (Cr) | SD ratio (Cr) | Mean Cramer-Rao | SD Cramer-Rao | |
| 0.41 | 0.10 | 11.36 | 2.78 | 0.42 | 0.10 | 10.96 | 2.11 | .48 |
| **Choline** | | | | | | | | |
| Pre-stressor | | | | Post-Stressor | | | | Paired t-test p-value |
| Mean ratio (Cr) | SD ratio (Cr) | Mean Cramer-Rao | SD Cramer-Rao | Mean ratio (Cr) | SD ratio (Cr) | Mean Cramer-Rao | SD Cramer-Rao | |
| .89 | .33 | 4.24 | .78 | .94 | .27 | 4.08 | .64 | .48 |
| **Glutamate** | | | | | | | | |
| Pre-stressor | | | | Post-Stressor | | | | Paired t-test p-value |
| Mean ratio (Cr) | SD ratio (Cr) | Mean Cramer-Rao | SD Cramer-Rao | Mean ratio (Cr) | SD ratio (Cr) | Mean Cramer-Rao | SD Cramer-Rao | |
| .96 | .16 | 5.80 | 1.47 | .97 | .17 | 5.92 | 2.20 | .76 |
| **Glutamine** | | | | | | | | |
| Pre-stressor | | | | Post-Stressor | | | | Paired t-test p-value |
| Mean ratio (Cr) | SD ratio (Cr) | Mean Cramer-Rao | SD Cramer-Rao | Mean ratio (Cr) | SD ratio (Cr) | Mean Cramer-Rao | SD Cramer-Rao | |
| .30 | .07 | 13.80 | 2.02 | .30 | .11 | 13.56 | 3.12 | .92 |
| **Myo-inositol** | | | | | | | | |
| Pre-stressor | | | | Post-Stressor | | | | Paired t-test p-value |
| Mean ratio (Cr) | SD ratio (Cr) | Mean Cramer-Rao | SD Cramer-Rao | Mean ratio (Cr) | SD ratio (Cr) | Mean Cramer-Rao | SD Cramer-Rao | |
| .63 | .21 | 5.12 | 1.36 | .70 | .30 | 5.48 | 1.42 | .27 |
| **N-Acetylaspartic Acid (NAA)** | | | | | | | | |
| Pre-stressor | | | | Post-Stressor | | | | Paired t-test p-value |
| Mean ratio (Cr) | SD ratio | Mean Cramer-Rao | SD Cramer-Rao | Mean ratio (Cr) | SD ratio | Mean Cramer-Rao | SD Cramer-Rao | |
| 1.10 | .13 | 2.84 | .75 | 1.16 | .14 | 3.00 | .96 | .04 |

**Supplementary Table 2b: Mean Cr ratios and Cramer-Rao standard deviations for MRS metabolites pre-stressor and post-stressor in the healthy control stress replication sample.**

| Aspartate | | | | | | | | |
|---|---|---|---|---|---|---|---|---|
| Pre-stressor | | | | Post-stressor | | | | Paired t-test p-value |
| Mean ratio (Cr) | SD ratio (Cr) | Mean Cramer-Rao | SD Cramer-Rao | Mean ratio (Cr) | SD ratio (Cr) | Mean Cramer-Rao | SD Cramer-Rao | |
| .49 | .10 | 9.09 | 2.49 | .48 | .09 | 9.50 | 1.95 | .40 |
| **Choline** | | | | | | | | |
| Pre-stressor | | | | Post-stressor | | | | Paired t-test p-value |
| Mean ratio (Cr) | SD ratio (Cr) | Mean Cramer-Rao | SD Cramer-Rao | Mean ratio (Cr) | SD ratio (Cr) | Mean Cramer-Rao | SD Cramer-Rao | |
| .72 | .11 | 3.86 | .64 | .75 | .13 | 4.27 | .70 | .32 |
| **Glutamate** | | | | | | | | |
| Pre-stressor | | | | Post-stressor | | | | Paired t-test p-value |
| Mean ratio (Cr) | SD ratio (Cr) | Mean Cramer-Rao | SD Cramer-Rao | Mean ratio (Cr) | SD ratio (Cr) | Mean Cramer-Rao | SD Cramer-Rao | |
| .96 | .21 | 5.64 | 2.06 | .97 | .26 | 5.36 | 2.08 | .88 |
| **Glutamine** | | | | | | | | |
| Pre-stressor | | | | Post-stressor | | | | Paired t-test p-value |
| Mean ratio (Cr) | SD ratio (Cr) | Mean Cramer-Rao | SD Cramer-Rao | Mean ratio (Cr) | SD ratio (Cr) | Mean Cramer-Rao | SD Cramer-Rao | |
| .22 | .04 | 12.14 | 2.59 | .21 | .05 | 12.86 | 3.26 | .72 |
| **Myo-inositol** | | | | | | | | |
| Pre-stressor | | | | Post-stressor | | | | Paired t-test p-value |
| Mean ratio (Cr) | SD ratio (Cr) | Mean Cramer-Rao | SD Cramer-Rao | Mean ratio (Cr) | SD ratio (Cr) | Mean Cramer-Rao | SD Cramer-Rao | |
| .80 | .14 | 4.45 | 1.26 | .80 | .09 | 4.00 | .98 | .85 |
| **N-Acetylaspartic Acid (NAA)** | | | | | | | | |
| Pre-Stressor | | | | Post-stressor | | | | Paired t-test p-value |
| Mean ratio (Cr) | SD ratio | Mean Cramer-Rao | SD Cramer-Rao | Mean ratio (Cr) | SD ratio | Mean Cramer-Rao | SD Cramer-Rao | |
| 1.06 | .23 | 3.05 | 1.40 | .98 | .32 | 3.41 | 1.74 | .05 |

**Supplementary Table 2c: Mean Cr ratios and Cramer-Rao standard deviations for MRS metabolites pre-no stress control (NSC) and post-NSC in the no stress control sample.**

| **Aspartate** | | | | | | | | |
|---|---|---|---|---|---|---|---|---|
| **Pre-NSC** | | | | **Post-NSC** | | | | Paired t-test p-value |
| Mean ratio (Cr) | SD ratio (Cr) | Mean Cramer-Rao | SD Cramer-Rao | Mean ratio (Cr) | SD ratio (Cr) | Mean Cramer-Rao | SD Cramer-Rao | |
| .42 | .09 | 11.83 | 3.34 | .42 | .09 | 11.06 | 4.14 | .78 |
| **Choline** | | | | | | | | |
| **Pre-NSC** | | | | **Post-NSC** | | | | Paired t-test p-value |
| Mean ratio (Cr) | SD ratio (Cr) | Mean Cramer-Rao | SD Cramer-Rao | Mean ratio (Cr) | SD ratio (Cr) | Mean Cramer-Rao | SD Cramer-Rao | |
| .71 | .07 | 4.56 | 1.42 | .72 | .11 | 4.61 | 1.38 | .92 |
| **Glutamate** | | | | | | | | |
| **Pre-NSC** | | | | **Post-NSC** | | | | Paired t-test p-value |
| Mean ratio (Cr) | SD ratio (Cr) | Mean Cramer-Rao | SD Cramer-Rao | Mean ratio (Cr) | SD ratio (Cr) | Mean Cramer-Rao | SD Cramer-Rao | |
| .98 | .21 | 5.83 | 2.07 | .94 | .16 | 6.56 | 1.62 | .30 |
| **Glutamine** | | | | | | | | |
| **Pre-NSC** | | | | **Post-NSC** | | | | Paired t-test p-value |
| Mean ratio (Cr) | SD ratio (Cr) | Mean Cramer-Rao | SD Cramer-Rao | Mean ratio (Cr) | SD ratio (Cr) | Mean Cramer-Rao | SD Cramer-Rao | |
| .25 | .08 | 13.89 | 2.89 | .24 | .09 | 13.67 | 3.51 | .52 |
| **Myo-inositol** | | | | | | | | |
| **Pre-NSC** | | | | **Post-NSC** | | | | Paired t-test p-value |
| Mean ratio (Cr) | SD ratio (Cr) | Mean Cramer-Rao | SD Cramer-Rao | Mean ratio (Cr) | SD ratio (Cr) | Mean Cramer-Rao | SD Cramer-Rao | |
| .84 | .15 | 4.61 | 1.14 | .81 | .16 | 4.50 | .99 | .45 |
| **N-Acetylaspartic Acid (NAA)** | | | | | | | | |
| **Pre-NSC** | | | | **Post-NSC** | | | | Paired t-test p-value |
| Mean ratio (Cr) | SD ratio | Mean Cramer-Rao | SD Cramer-Rao | Mean ratio (Cr) | SD ratio | Mean Cramer-Rao | SD Cramer-Rao | |
| 1.03 | .17 | 3.33 | 1.57 | 1.00 | .18 | 3.61 | 1.29 | .24 |

**Supplementary Table 2d: Mean Cr ratios and Cramer-Rao standard deviations for MRS metabolites for pre-stressor and post-stressor acquisitions in the major depressive disorder sample.**

| Aspartate | | | | | | | | |
|---|---|---|---|---|---|---|---|---|
| Pre-stressor | | | | Post-stressor | | | | Paired t-test p-value |
| Mean ratio (Cr) | SD ratio (Cr) | Mean Cramer-Rao | SD Cramer-Rao | Mean ratio (Cr) | SD ratio (Cr) | Mean Cramer-Rao | SD Cramer-Rao | |
| .47 | .11 | 10.87 | 2.32 | .48 | .11 | 10.70 | 2.53 | .47 |

| Choline | | | | | | | | |
|---|---|---|---|---|---|---|---|---|
| Pre-stressor | | | | Post-stressor | | | | Paired t-test p-value |
| Mean ratio (Cr) | SD ratio (Cr) | Mean Cramer-Rao | SD Cramer-Rao | Mean ratio (Cr) | SD ratio (Cr) | Mean Cramer-Rao | SD Cramer-Rao | |
| .71 | .10 | 4.22 | .80 | .80 | .22 | 4.22 | .95 | .06 |

| Glutamate | | | | | | | | |
|---|---|---|---|---|---|---|---|---|
| Pre-stressor | | | | Post-stressor | | | | Paired t-test p-value |
| Mean ratio (Cr) | SD ratio (Cr) | Mean Cramer-Rao | SD Cramer-Rao | Mean ratio (Cr) | SD ratio (Cr) | Mean Cramer-Rao | SD Cramer-Rao | |
| .99 | .16 | 5.61 | 1.88 | .98 | .26 | 6.43 | 2.13 | .81 |

| Glutamine | | | | | | | | |
|---|---|---|---|---|---|---|---|---|
| Pre-stressor | | | | Post-stressor | | | | Paired t-test p-value |
| Mean ratio (Cr) | SD ratio (Cr) | Mean Cramer-Rao | SD Cramer-Rao | Mean ratio (Cr) | SD ratio (Cr) | Mean Cramer-Rao | SD Cramer-Rao | |
| .24 | .06 | 14.00 | 2.43 | .25 | .08 | 13.61 | 3.30 | .46 |

| Myo-inositol | | | | | | | | |
|---|---|---|---|---|---|---|---|---|
| Pre-stressor | | | | Post-stressor | | | | Paired t-test p-value |
| Mean ratio (Cr) | SD ratio (Cr) | Mean Cramer-Rao | SD Cramer-Rao | Mean ratio (Cr) | SD ratio (Cr) | Mean Cramer-Rao | SD Cramer-Rao | |
| .79 | .11 | 4.61 | 1.12 | .77 | .18 | 4.57 | 1.04 | .71 |

| N-Acetylaspartic Acid (NAA) | | | | | | | | |
|---|---|---|---|---|---|---|---|---|
| Pre-stressor | | | | Post-stressor | | | | Paired t-test p-value |
| Mean ratio (Cr) | SD ratio | Mean Cramer-Rao | SD Cramer-Rao | Mean ratio (Cr) | SD ratio | Mean Cramer-Rao | SD Cramer-Rao | |
| 1.02 | .16 | 3.00 | 1.09 | .95 | .19 | 3.87 | 1.14 | .05 |

**Supplementary Table 3**: Hierarchical regression models predicting %ΔGlu and %ΔGlx in healthy control samples.

Acute-perceived stress interaction in healthy controls (Glu)

| Model 1 | Predictor | B | SE | Beta | *t* | *p* | adj-$r^2$ | *F* | *p* | $r^2$ change | *F change* | *Sig F Change* |
|---|---|---|---|---|---|---|---|---|---|---|---|---|
| Step 1 | Acute Stress | 0.029 | 0.072 | 0.049 | 0.399 | 0.692 | 0.08 | 3.81 | 0.027 | 0.11 | 3.81 | 0.027 |
| | PSS | -0.018 | 0.007 | -0.319 | -2.619 | 0.011 | | | | | | |

| Model 2 | Predictor | B | SE | Beta | *t* | *p* | adj-$r^2$ | *F* | *p* | $r^2$ change | *F change* | *Sig F Change* |
|---|---|---|---|---|---|---|---|---|---|---|---|---|
| Step 1 | Acute Stress | -0.008 | 0.071 | -0.013 | -0.111 | 0.912 | 0.15 | 4.85 | 0.004 | 0.08 | 6.27 | 0.015 |
| | PSS | -0.013 | 0.007 | -0.240 | -1.979 | 0.052 | | | | | | |
| Step 2 | Age | 0.012 | 0.005 | 0.308 | 2.503 | 0.015 | | | | | | |
| | PSS x Acute Stress | | | | -2.030 | 0.047 | | | | | | |
| | Sex | | | | 0.700 | 0.486 | | | | | | |
| | Study Site | | | | -0.643 | 0.522 | | | | | | |

| Model 3 | Predictor | B | SE | Beta | *t* | *p* | adj-$r^2$ | *F* | *p* | $r^2$ change | *F change* | *Sig F Change* |
|---|---|---|---|---|---|---|---|---|---|---|---|---|
| Step 1 | Acute Stress | 0.011 | 0.070 | 0.019 | 0.158 | 0.875 | 0.19 | 4.85 | 0.002 | 0.05 | 4.12 | 0.047 |
| | PSS | 0.002 | 0.010 | 0.043 | 0.237 | 0.813 | | | | | | |
| Step 2 | Age | 0.011 | 0.005 | 0.289 | 2.399 | 0.020 | | | | | | |
| | PSS x Acute Stress | -0.026 | 0.013 | -0.365 | -2.030 | 0.047 | | | | | | |
| | Sex | | | | 0.086 | 0.932 | | | | | | |
| | Study Site | | | | -0.809 | 0.422 | | | | | | |

Acute-perceived stress interaction in healthy controls (Glx)

| Model 1 | Predictor | B | SE | Beta | *t* | *p* | adj-$r^2$ | *F* | *p* | $r^2$ change | *F change* | *Sig F Change* |
|---|---|---|---|---|---|---|---|---|---|---|---|---|
| Step 1 | Acute Stress | 0.022 | 0.067 | 0.038 | 0.323 | 0.748 | 0.12 | 5.32 | 0.007 | 0.15 | 5.32 | 0.007 |
| | PSS | -0.020 | 0.006 | -0.374 | -3.136 | 0.003 | | | | | | |

| Model 2 | Predictor | B | SE | Beta | *t* | *p* | adj-$r^2$ | *F* | *p* | $r^2$ change | *F change* | *Sig F Change* |
|---|---|---|---|---|---|---|---|---|---|---|---|---|
| Step 1 | Acute Stress | -0.007 | 0.067 | -0.012 | -0.098 | 0.922 | 0.16 | 5.11 | 0.003 | 0.06 | 4.16 | 0.046 |
| | PSS | -0.016 | 0.006 | -0.310 | -2.571 | 0.013 | | | | | | |
| Step 2 | Age | 0.009 | 0.005 | 0.250 | 2.041 | 0.046 | | | | | | |
| | PSS x Acute Stress | | | | -1.802 | 0.077 | | | | | | |
| | Sex | | | | 0.553 | 0.583 | | | | | | |
| | Study Site | | | | -0.677 | 0.501 | | | | | | |

*Note*. Stress condition (acute stress or no stress control; "acute stress") and PSS were entered in the first block and all other predictors were included in the second block with stepwise selection. Model coefficients (adjusted $r^2$, $F$) are based on included variables, while $t$ and $p$ are provided for excluded variables. SE = Standard Error; PSS = Perceived Stress Scale.

**Supplementary Table 4**: Hierarchical regression models predicting %ΔGlu and %ΔGlx in all stress samples.

Diagnostic Group x PSS interaction in individuals who completed the acute stress manipulation (Glu)

| Model 1 | Predictor | B | SE | Beta | *t* | *p* | adj-$r^2$ | *F* | *p* | $r^2$ change | *F change* | *Sig F Change* |
|---|---|---|---|---|---|---|---|---|---|---|---|---|
| Step 1 | Diagnostic Group | 0.203 | 0.152 | 0.329 | 1.337 | 0.186 | 0.021 | 1.727 | 0.186 | 0.049 | 1.727 | 0.186 |
| | PSS | -0.013 | 0.007 | -0.442 | -1.795 | 0.077 | | | | | | |

| Model 2 | Predictor | B | SE | Beta | *t* | *p* | adj-$r^2$ | *F* | *p* | $r^2$ change | *F change* | *Sig F Change* |
|---|---|---|---|---|---|---|---|---|---|---|---|---|
| Step 1 | Diagnostic Group | 0.056 | 0.157 | 0.090 | 0.354 | 0.724 | 0.096 | 3.431 | 0.022 | 0.086 | 6.552 | 0.013 |
| | PSS | -0.029 | 0.009 | -0.986 | -3.100 | 0.003 | | | | | | |
| Step 2 | PSS x Diagnostic Group | 0.036 | 0.014 | 0.816 | 2.560 | 0.013 | | | | | | |
| | Sex | | | | -0.225 | 0.823 | | | | | | |
| | Age | | | | 1.032 | 0.306 | | | | | | |
| | Study Site | | | | -0.435 | 0.665 | | | | | | |

Diagnostic Group x PSS interaction in individuals who completed the acute stress manipulation (Glx)

| Model 1 | Predictor | B | SE | Beta | *t* | *p* | adj-$r^2$ | *F* | *p* | $r^2$ change | *F change* | *Sig F Change* |
|---|---|---|---|---|---|---|---|---|---|---|---|---|
| Step 1 | Diagnostic Group | 0.199 | 0.142 | 0.346 | 1.401 | 0.166 | 0.013 | 1.472 | 0.237 | 0.042 | 1.472 | 0.237 |
| | PSS | -0.012 | 0.007 | -0.421 | -1.705 | 0.093 | | | | | | |

| Model 2 | Predictor | B | SE | Beta | *t* | *p* | adj-$r^2$ | *F* | *p* | $r^2$ change | *F change* | *Sig F Change* |
|---|---|---|---|---|---|---|---|---|---|---|---|---|
| Step 1 | Diagnostic Group | 0.040 | 0.145 | 0.069 | 0.275 | 0.784 | 0.120 | 4.122 | 0.010 | 0.166 | 9.068 | 0.004 |
| | PSS | -0.029 | 0.009 | -1.053 | -3.356 | 0.001 | | | | | | |
| Step 2 | PSS x Diagnostic Group | 0.039 | 0.013 | 0.948 | 3.011 | 0.004 | | | | | | |
| | Sex | | | | -0.302 | 0.764 | | | | | | |
| | Age | | | | -0.509 | 0.613 | | | | | | |
| | Study Site | | | | 0.742 | 0.461 | | | | | | |

*Note*. Diagnostic group and PSS were entered in the first block and all other predictors were included in the second block with stepwise selection. Model coefficients (adjusted $r^2$, *F*) are based on included variables, while *t* and *p* are provided for excluded variables. SE = Standard Error; PSS = Perceived Stress Scale.

**Supplementary Table 5**: Linear vs quadratic effects of PSS on %∆Glu and %∆Glx after acute stress.

Quadratic Effect of PSS (Glu)

| Model 1 | Predictor | B | SE | Beta | *t* | *p* | adj-$r^2$ | *F* | *p* | $r^2$ change | *F change* | *Sig F Change* |
|---|---|---|---|---|---|---|---|---|---|---|---|---|
| | PSS | -0.005 | 0.004 | -0.154 | -1.284 | 0.204 | 0.009 | 1.648 | 0.204 | 0.024 | 1.648 | 0.204 |

| Model 2 | Predictor | B | SE | Beta | *t* | *p* | adj-$r^2$ | *F* | *p* | $r^2$ change | *F change* | *Sig F Change* |
|---|---|---|---|---|---|---|---|---|---|---|---|---|
| | PSS | -0.010 | 0.004 | -0.347 | -2.426 | 0.018 | 0.069 | 3.553 | 0.034 | 0.072 | 5.352 | 0.024 |
| | PSS Squared | 0.001 | 0.000 | 0.331 | 2.313 | 0.024 | | | | | | |

Quadratic Effect of PSS (Glx)

| Model 1 | Predictor | B | SE | Beta | *t* | *p* | adj-$r^2$ | *F* | *p* | $r^2$ change | *F change* | *Sig F Change* |
|---|---|---|---|---|---|---|---|---|---|---|---|---|
| | PSS | -0.003 | 0.003 | -0.118 | -0.983 | 0.329 | 0.000 | 0.966 | 0.328 | 0.014 | 0.966 | 0.329 |

| Model 2 | Predictor | B | SE | Beta | *t* | *p* | adj-$r^2$ | *F* | *p* | $r^2$ change | *F change* | *Sig F Change* |
|---|---|---|---|---|---|---|---|---|---|---|---|---|
| | PSS | -0.009 | 0.004 | -0.342 | -2.408 | 0.019 | 0.084 | 4.160 | 0.020 | 0.096 | 7.264 | 0.009 |
| | PSS Squared | 0.001 | 0.000 | 0.383 | 2.695 | 0.009 | | | | | | |

*Note*. SE = Standard Error; PSS = Perceived Stress Scale.

**Supplementary Table 6:** Hierarchical regression models testing stress by age and stress by sex interactions on %ΔGlu and %ΔGlx in healthy controls.

Sex interaction with stress in healthy control stress samples (Glu)

| Model 1 | **Predictor** | **B** | **SE** | **Beta** | ***t*** | ***p*** | **adj-$r^2$** | ***F*** | ***p*** |
|---|---|---|---|---|---|---|---|---|---|
| Step 1 | PSS | -0.029 | 0.010 | -0.424 | -2.985 | 0.005 | 0.151 | 5.101 | 0.010 |
| | Sex | 0.018 | 0.089 | 0.029 | 0.201 | 0.842 | | | |
| Step 2 | Sex x PSS | | | | -0.934 | 0.355 | | | |
| | Study Site | | | | -0.483 | 0.632 | | | |

Age interaction with stress in healthy control stress samples (Glu)

| Model 1 | **Predictor** | **B** | **SE** | **Beta** | ***t*** | ***p*** | **adj-$r^2$** | ***F*** | ***p*** |
|---|---|---|---|---|---|---|---|---|---|
| Step 1 | PSS | -0.024 | 0.009 | -0.360 | -2.668 | 0.011 | 0.223 | 7.588 | 0.001 |
| | Age | 0.011 | 0.005 | 0.273 | 2.020 | 0.049 | | | |
| Step 2 | Age x PSS | | | | 0.835 | 0.408 | | | |
| | Study Site | | | | -0.743 | 0.462 | | | |

Sex interaction with stress in healthy control stress samples (Glx)

| Model 1 | **Predictor** | **B** | **SE** | **Beta** | ***t*** | ***p*** | **adj-$r^2$** | ***F*** | ***p*** |
|---|---|---|---|---|---|---|---|---|---|
| Step 1 | PSS | -0.028 | 0.009 | -0.452 | -3.238 | 0.002 | 0.181 | 6.075 | 0.005 |
| | Sex | 0.022 | 0.081 | 0.038 | 0.272 | 0.787 | | | |
| Step 2 | Sex x PSS | | | | -1.621 | 0.112 | | | |
| | Study Site | | | | -0.574 | 0.569 | | | |

Age interaction with stress in healthy control stress samples (Glx)

| Model 1 | **Predictor** | **B** | **SE** | **Beta** | ***t*** | ***p*** | **adj-$r^2$** | ***F*** | ***p*** |
|---|---|---|---|---|---|---|---|---|---|
| Step 1 | PSS | -0.025 | 0.008 | -0.402 | -2.996 | 0.004 | 0.231 | 7.916 | 0.001 |
| | Age | 0.009 | 0.005 | 0.231 | 1.722 | 0.092 | | | |
| Step 2 | Age x PSS | | | | 0.974 | 0.335 | | | |
| | Study Site | | | | -0.780 | 0.440 | | | |

*Note*. For each model, PSS and the main demographic variable of interest were entered in the first block and interaction terms were included in the second block with stepwise selection. Model coefficients (adjusted $r^2$, $F$) are based on included variables, while $t$ and $p$ are provided for excluded variables. SE = Standard Error; PSS = Perceived Stress Scale.

**Supplementary Table 7:** Hierarchical regression models testing effects of birth control on %ΔGlu and %ΔGlx in healthy control female participants.

Birth control interaction with PSS in female healthy controls following acute stress (Glu)

| Model 1 | Predictor | B | SE | Beta | *t* | *p* | adj-$r^2$ | *F* | *p* |
|---|---|---|---|---|---|---|---|---|---|
| Step 1 | PSS | -0.034 | 0.013 | -0.443 | -2.648 | 0.013 | 0.161 | 3.971 | 0.030 |
| | Birth Control (No/Yes) | -0.049 | 0.107 | -0.077 | -0.463 | 0.647 | | | |
| Step 2 | Birth Control x PSS | | | | 0.526 | 0.603 | | | |
| | Study Site | | | | -0.417 | 0.680 | | | |

Birth control interaction with PSS in female healthy controls following acute stress (Glx)

| Model 1 | Predictor | B | SE | Beta | *t* | *p* | adj-$r^2$ | *F* | *p* |
|---|---|---|---|---|---|---|---|---|---|
| Step 1 | PSS | -0.037 | 0.011 | -0.535 | -3.438 | 0.002 | 0.274 | 6.841 | 0.004 |
| | Birth Control (No/Yes) | -0.064 | 0.090 | -0.111 | -0.711 | 0.482 | | | |
| Step 2 | Birth Control x PSS | | | | 0.639 | 0.528 | | | |
| | Study Site | | | | -0.577 | 0.569 | | | |

*Note*. PSS and birth control (no/yes) were entered in the first block and interaction terms were included in the second block with stepwise selection. Model coefficients (adjusted $r^2$, $F$) are based on included variables, while $t$ and $p$ are provided for excluded variables. SE = Standard Error; PSS = Perceived Stress Scale.

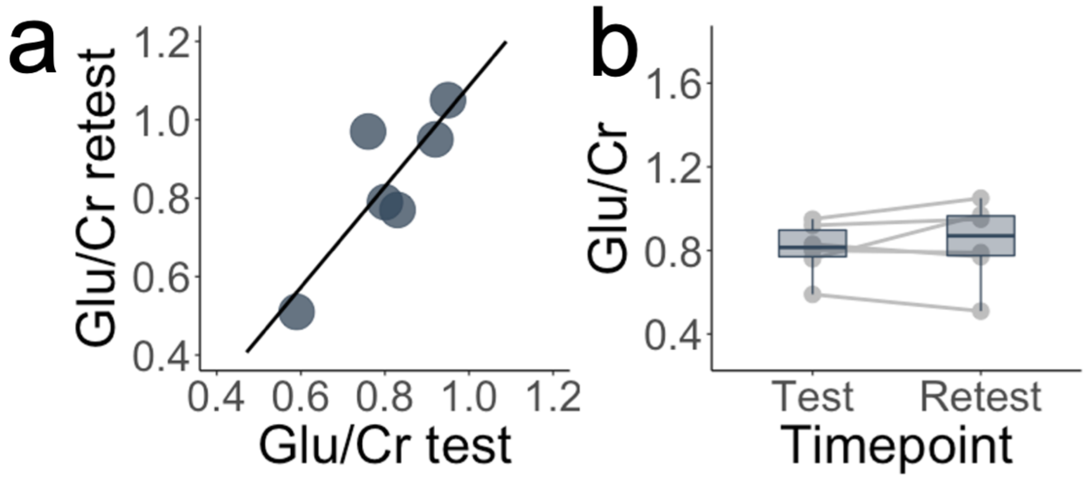


**<u>Supplementary Figure 1</u>:** Test-retest of MRS acquisition sequence at Emory University. **a)** Scatter plot for test-retest MRS data collected in six subjects during the same scanning session with no stress manipulation in between MRS acquisitions (Intra-class correlation coefficient = 0.89, $p$ = .017). These data were collected at Emory University using the same Siemens Tim Trio and identical MRS protocol. **b)** Box plot showing reliability of test-retest data.

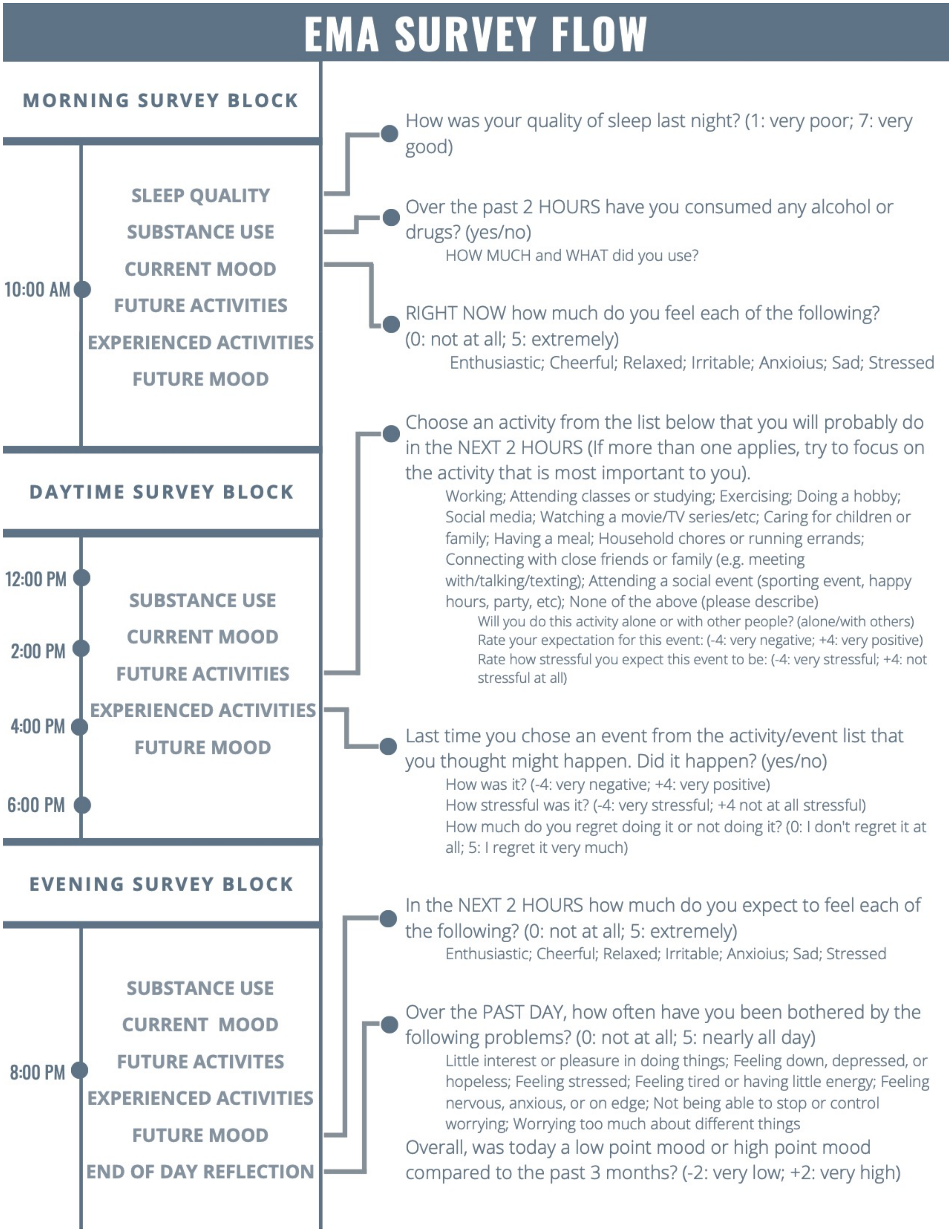


**<u>Supplementary Figure 2:</u>** Ecological momentary assessment (EMA) items and survey flow. Individual questions (left) were sent 6 times per day, every other day, according to survey flow (right). Median start time was 10am and ranged from 6am to 12pm, depending on the participant's schedule.

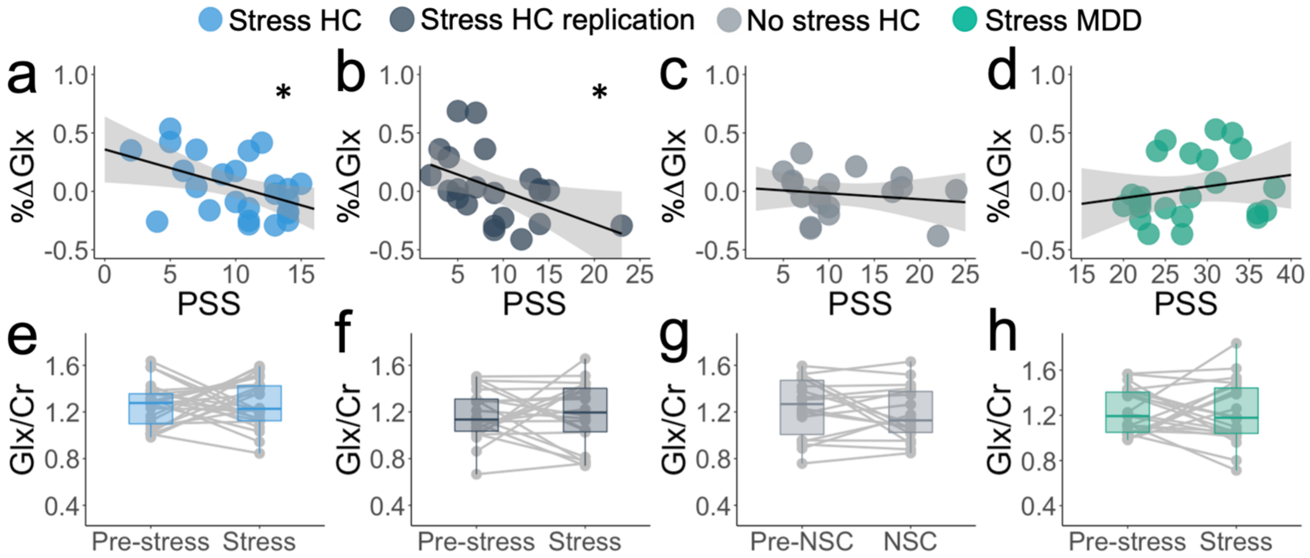


**Supplementary Figure 3:** Changes in mPFC Glx (Glutamate + Glutamine) in response to acute and perceived stress **a)** Association between perceived stress (PSS scores) and percent change in MRS Glx signal (%ΔGlx; $r_s$ = -.401, *p* = .047) in healthy control stress sample. **b)** Association between perceived stress (PSS scores) and percent change in MRS Glx signal (%ΔGlx; $r_s$ = -.465, *p* = .029) in the healthy control stress replication sample. **c)** Association between perceived stress (PSS scores) and percent change in MRS Glx signal (%ΔGlx; $r_s$ = -.214, *p* = .395) in no stress control sample. **d)** Association between perceived stress (PSS scores) and percent change in MRS Glx signal (%ΔGlx; $r_s$ = .189, *p* = .387) in participants with major depressive disorder. Shaded area on a-d represents 95% confidence interval, **p*<.05. **e-h.** Glx/Cr ratios before and after MAST in healthy control stress sample (e), healthy control stress replication (f), no stress control (g), and subjects with major depressive disorder (h).

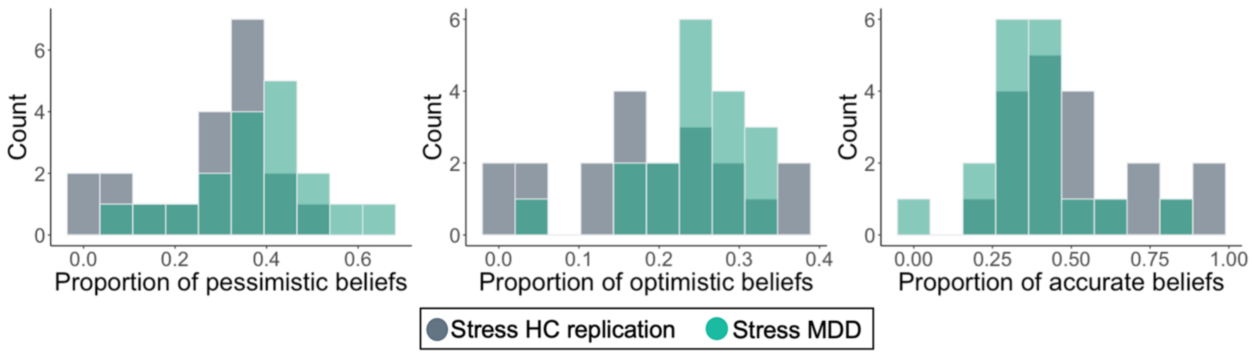


**Supplementary Figure 4:** Relative frequencies of accurate and inaccurate expectations from ecological momentary assessment (EMA) data. The proportion of observations of each type, relative to total observations, were computed for each participant. Histograms represent the distribution of proportions for pessimistic beliefs (left), optimistic beliefs (middle), and accurate beliefs (right).
